# Microplastics are detected in bull and dog sperm and polystyrene microparticles impair sperm fertilization

**DOI:** 10.1101/2023.12.15.571802

**Authors:** N. Grechi, G.A. Ferronato, S. Devkota, M.A.M.M. Ferraz

## Abstract

The alarming increase in global infertility rates has coincided with the pervasive accumulation of microplastics (MPs) resulting from the poor management of plastic waste. This concerning trend is particularly troubling because only 10% of male infertility cases can be attributed to identifiable causes, leaving a significant knowledge gap in our understanding of their underlying factors. To bridge this critical gap, it is important to explore the connection between the accumulation of MPs and the observed decline in male fertility. Here, the presence of microplastics in reproductive fluids from bulls and dogs was assessed and used as baseline concentrations for bull sperm exposure. Bovine epididymal sperm (ES) presented a mean of 72.5 MP particles mL^-1^ (0.3691 μg mL^−1^) while canine seminal plasma had an average of 35.4 MP particles mL^-1^ (0.0066 μg mL^−1^). Bovine sperm was exposed to three different concentrations of a mixture of 1.1, 0.5, and 0.3 µm polystyrene (PS) beads: (1) 0.7 μg mL^−1^, blood concentration of PS in cows (bPS); (2) 0.37 μg mL^−1^, concentration of total MPs in ES (esMP); and (3) 0.026 μg mL^−1^, concentration of PS in ES (esPS). All sperm samples incubated with PS exhibited reduced motility compared with the CT at 0.5 h. However, PS exposure did not affect acrosome or induced oxidative stress. When used for *in vitro* fertilization, the sperm exposed to PS had decreased blastocyst rates, in addition to inducing ROS formation and apoptosis on resulting embryos. By employing realistic exposure concentrations, this research sought to shed light on the comprehensive impact of MPs on bovine sperm and the quality of resulting embryos, providing the first evidence of MPs in bovine and dog sperm and demonstrating the detrimental effect of PS MPs on sperm motility and functionality.

**Graphical Abstract:** 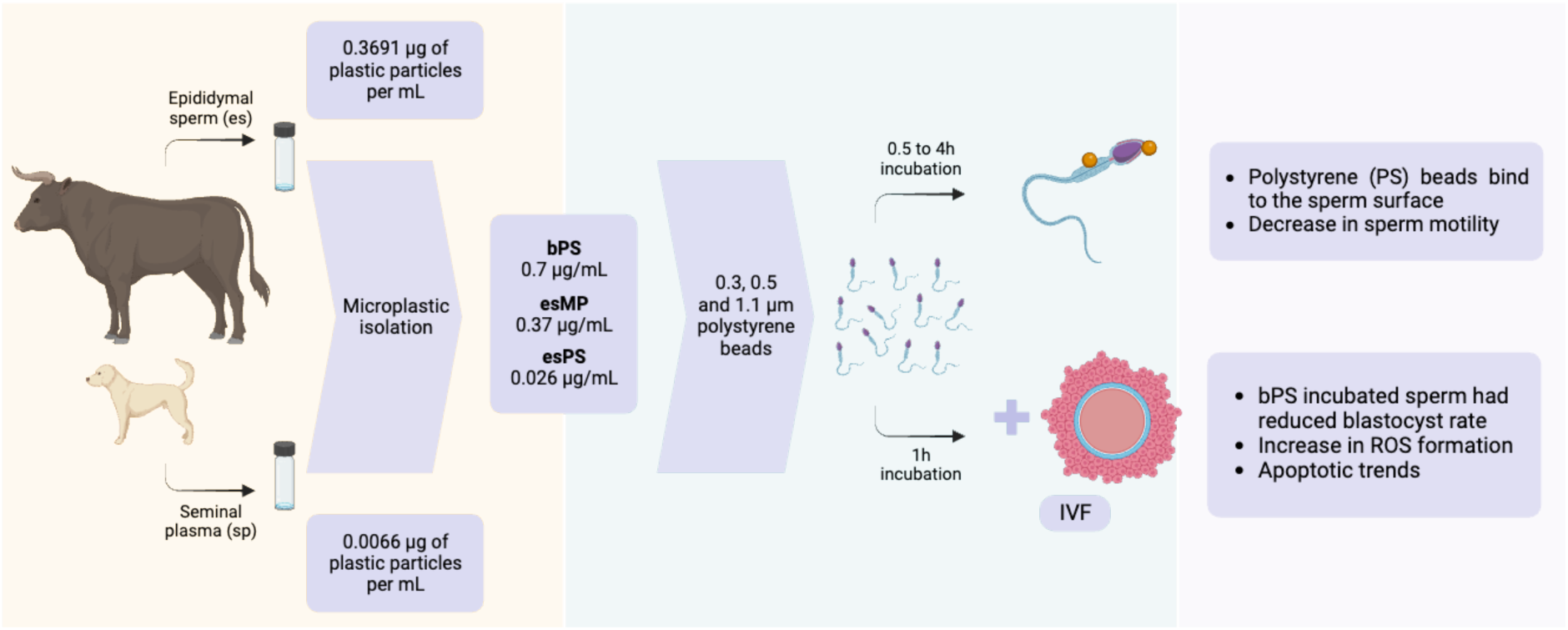

**Highlights:**

1. Microplastics were found across the most diverse range of environments and their presence have been shown to affect reproductive parameters within different species.
2. True-to-life concentrations of exposure were used to assess the potential effects of polystyrene in sperm parameters and fertilization.
3. Polystyrene microplastics attach to sperm and decrease motility, also reducing sperm functionality as seen by decreased blastocyst rate and increased oxidative stress in embryos.

## 1. Introduction

Since their first synthesis, plastics have been part of several aspects of modern life, with diverse applications in human activities. The inadequate management practices surrounding plastics have resulted in them becoming a prominent source of pollution, contributing significantly to public health and environmental problems (Velis et al., 2023). As a consequence, there has been a distressing accumulation of microplastics (MPs)(Semmouri et al., 2023). While the existence of MPs was initially identified in 1970, it was not until 2004 that they were referred to as plastic fragments measuring a few micrometres in diameter (Thompson et al., 2004). Only within the past decade have researchers begun to pay substantial attention to the concerning accumulation of these particles (Napper and Thompson, 2020). Presently, MPs have been ubiquitously detected in the air, water, soil, and even the food we consume, posing an imminent threat to human, animal, and environmental health (Pironti et al., 2021). Of concern, MPs were also detected in a range of human and animal samples, including lung (Amato-Lourenço et al., 2021; Jenner et al., 2022), placenta (Braun et al., 2021; Ragusa et al., 2021), testis (Zhao et al., 2023a), feces (Liu et al., 2022; Luqman et al., 2021; Schwabl et al., 2019), sputum (Huang et al., 2022), blood (Carrington, 2022), milk (Liu et al., 2022), amniotic fluid (Halfar et al., 2023), follicular fluid (Grechi et al., 2023) and semen (Montano et al., 2023; Zhao et al., 2023a).

Simultaneously, there has been a substantial increase in infertility rates across both developed and developing nations in recent decades (Borumandnia et al., 2022). The impact of this condition is far-reaching, affecting one in every six individuals of reproductive age worldwide (World Health Organization (WHO), 2018). Among all these known cases, the male factor is responsible for 30% of them alone and contributes to 20% of the combined infertility cases (Agarwal et al., 2020). It is important to highlight that the known causes of male infertility only account for a mere 10% of the cases, suggesting a vast number of underlying factors that remain unidentified (Ávila et al., 2022). As emphasized by Levine et al., there was a worldwide decline of 51.6% in sperm concentration from 1973 to 2018, along with an overall decrease of 62.3% in total sperm count (Levine et al., 2023). While the precise causes are still to be entirely elucidated, potential factors include the exposure to environmental pollutants (Levine et al., 2023). Recent research has highlighted that the oral exposure of mice to polystyrene results in testicular and sperm toxicity, thereby affecting male fertility through the induction of oxidative stress (Xu et al., 2023). In humans, the presence of MPs in semen and testis had already been described, where the particles could have induced a disruption of the blood-testis-barrier and increased its permeability, making their way into the seminal fluid (Montano et al., 2023; Zhao et al., 2023a). Despite the observed impacts of MPs on sperm, there remains limited understanding of their potential implications for fertilization. Research conducted *in ovo* has demonstrated that polystyrene-exposed fertilized chicken eggs exhibit developmental delays, heightened mortality, and the occurrence of teratogenic effects alongside organ malformations (Hou et al., 2022). Similar findings have emerged from studies involving zebrafish, which offer a more comprehensive understanding of the consequences of MPs exposure on offspring, encompassing neurotoxicity and disruptions to the endocrine system (Torres-Ruiz et al., 2023). These studies have expanded our understanding of how MPs can interfere with embryo development, yet they do not elucidate how the impact on sperm contributes to such outcomes.

While these studies are crucial for uncovering the mechanisms through which continuous exposure to various types of MPs might affect fertility, they lack biological accuracy. The majority of these experiments involved feeding animals or exposing cells to MPs sometimes at levels thousands of times higher than what they would naturally encounter in their environment (Mills et al., 2023). Therefore, we investigated the presence of MPs in bovine epididymal sperm and dog seminal plasma and used it as baseline data for investigating the effects of MPs on bovine male gamete *in vitro*. We then investigated the potential impairments these sperm could impose on fertilization and embryo development.

## 2. Materials and methods

### 2.1 MP contamination prevention

In order to avoid any potential contamination from airborne or material MPs, all procedures were carried out inside a laminar flow hood, as we have demonstrated it to have the least environmental MP contamination during sample preparation (Noonan et al., 2023). Glass materials were used whenever possible to replace flasks and other apparatuses. To prevent material contamination of MPs, all materials and equipment utilized underwent a thorough three times rinsing process using ultrapure water that had been previously filtered using a 0.1 μm membrane (Merck Isopore). Furthermore, all reagents and water employed in the described protocols were filtered using a 0.1 μm membrane before being used. All samples and reagents were covered with aluminium foil during all procedures.

### 2.2 Microplastic polystyrene beads

Polystyrene beads (Latex Microsphere Suspension) with sizes of 0.31, 0.50, and 1.1 μm were purchased from Thermo Fisher Scientific. The beads were in water solution, at 10% w/w concentration. For the exposure assessments, those were diluted to the requirements of each experiment.

### 2.3 Sample collection

The seminal plasma samples from dogs were donated by the Clinic of Small Animal Surgery and Reproduction from the Ludwig-Maximilians-Universität München, samples were collected during normal clinical routine (for sperm banking) by digital manipulation (Seager et al., 1975), together with water controls from each step of the collection which was performed directly in glass vials. Bovine testicles, from animals between 19 and 22 months old, were obtained from a local slaughterhouse and immediately transported to the laboratory at room temperature (RT). Upon arrival, the testes were rinsed with distilled water and ethanol 70%, dried, and the tunica *vaginalis* was cut open to expose the testis and the epididymis. Half of the epididymis body and tail were dissected, external blood vessels removed, and the piece put on a glass petri dish on a warming plate (37°C). Filtered PBS (5 mL) was added, and several cuts made throughout the whole organ, releasing the epididymal sperm and fluids. After 5 min, several washes with PBS were performed to detach the epididymal sperm and fluids from the dish, and the liquid was transferred to a pre-rinsed glass vial. A blank control (PBS) was also collected according to the same protocol. All the samples were frozen (-20°C) until isolation.

### 2.4 Microplastic isolation from bovine epididymal sperm and canine seminal plasma

Canine seminal plasma (N=2) and bovine epididymal sperm (N=3) were used for microplastic isolation using a protocol we developed previously (Grechi et al., 2023). On the first day of isolation, 2 mL of each sample were put in an Erlenmeyer together with 50 mL of KOH 10% and incubated on a shaker at 60°C and 250 rpm for 24 h (1:25, sample:KOH 10%). Following the first 24 h, 10 mL of NaClO 47% was added to each of the samples, to reach a final concentration of 7.5%, and kept on a shaker at 60°C and 250 rpm, for another 24 h. After the second day of digestion, all the samples were filtered through a 47 mm polytetrafluoroethylene polymer (PTFE) membrane with a 0.45 μm pore (Merck Omnipore^TM^) and rinsed three times with filtered ultrapure water. Then, the membranes containing the particles were submersed in 50 mL of HNO_3_ 20%, put in an ultrasonic bath in sweep function with 100% power and 45 kHz frequency for 15 min, to transfer particles from the membrane to the solution, and incubated for 30 min at 40°C and 250 rpm. The membranes were then removed, and the samples were incubated for 24 h at 40°C and 250 rpm. On the final day, the samples were filtered through a 13 mm PTFE membrane with 0.45 μm pore and washed three times with filtered ultrapure water before being mounted on a glass slide for posterior microscopy and spectroscopy analysis. To avoid contamination during transportation and storage, the slides were covered until imaging. The above-mentioned blank controls from the seminal plasma collection (N = 2) and from epididymal sperm collection (N = 3) were processed according to the same protocol, together with each of the samples from the collection to the microscopy analysis.

### 2.5 Confocal Raman analysis of isolated particles

The microscopy and spectroscopy analysis involved the use of a confocal Raman microscope (alpha300 R, WITec, Germany) equipped with a spectrometer (UHTS 300, WITec, Germany) and a 532 nm laser. The microscope was operated using Control Five software, and the acquired data were processed using Project Five software, which enabled background noise and cosmic radiation signal compensation. Initially, a whole membrane image was generated by focusing a x10 NA 0.25 Zeiss objective and employing the Area Stitching option. From the whole membrane image, a specific field was selected and imaged again using a x50 NA 0.75 Zeiss objective, along with the Area Stitching option and Z-stack auto-focus. The Particle Scout software was used to identify the particles for which Raman spectra were obtained. Raman spectra measurements were conducted using a x100 DIC NA 0.9 Zeiss objective, with a laser power of 10 mV, accumulation of 4, and an integration time of 0.45 seconds. Spectra autofocus was employed during the measurements. After acquiring the spectra, the particles were compared against the S.T. Japan database (S.T. Japan Europe GmbH, Germany) and the SloPP MPs Raman library (Munno et al., 2020) using the True Match-Integrated Raman Spectra Database Management software. Matches with a hit quality index (HQI) greater than 75% were considered for further analysis. Particles made of Poly(tetrafluoroethylene-co-perfluoro-(alkyl vinyl ether)) and Perfluoroalkoxy alkane (PFA) were excluded from the analysis as they are components of the pore membrane used for imaging. After the process of matching and excluding membrane particles, the quantification of microplastics was fine-tuned by subtracting the count of particles found in the water control samples from the count of particles in the sample of interest, with this adjustment performed individually for each specific polymer detected.

As previously described (Grechi et al., 2023), all the particles that were identified during the analysis were categorized into seven different classes: 1) non-plastic related particles (NP); 2) plastic polymers (PP); 3) plasticizers (PZ); 4) pigments (PG); 5) coatings, solvents, or fillers (CSF); 6) fibers (FB); and 7) unknown particles (UK). To determine the weight of each particle, the following formula was used:

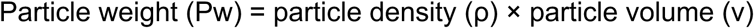

### 2.6 Microplastic concentrations for exposure assessment

Three concentrations of microplastics were selected based on previous literature and the analysis conducted by the research group. These concentrations were determined using the weight of polystyrene (density 1.05 g mL^-1^) and the equation described above. A stock solution with a concentration of 10 mg mL^-1^ was prepared, containing equal weights of polystyrene beads with diameters of 0.3 μm, 0.5 μm, and 1.1 μm. Subsequent dilutions, to which the sperm samples were exposed, were prepared using this stock solution. The first concentration used in the experiments was 0.7 μg mL^-1^ (bPS), which was derived from the total amount of polystyrene found in cow blood, as reported by (Veen et al., 2022). The second concentration, 0.369 μg mL^-1^ (esMP), was obtained from the total microplastics identified in the epididymal sperm samples from bulls. Finally, the third concentration of 0.0257 μg mL^-1^ (esPS) was obtained from the polystyrene particles detected in the epididymal sperm samples from bulls.

### 2.7 Sperm incubation with MPs

Frozen bull sperm were thawed at 37°C for 30 s, the content of the straws was added to 2 mL of semen preparation medium (BO-SemenPrep™, IVF Bioscience, UK), centrifuged for 5 min at 330xg, the pellet resuspended with 2 mL of the same media, and again centrifuged (330xg, 5 min). The supernatant was removed, and the remaining pellet was resuspended with 500 μL of Tyrode’s medium supplemented with 2 mM sodium bicarbonate, 10 mM lactate, 1 mM pyruvate, 6 mg mL^-1^ fatty acid–free bovine serum albumin (BSA), 100 U mL^-1^ penicillin and 100 μg mL^-1^ streptomycin (RD media). The after-thaw motility and concentration were assessed with a mobile computer-assisted sperm analyzer iSperm® mCASA (Aidmics Biotechnology, Taipei, Taiwan).

As a pilot study, 5x10^6^ sperm mL^-1^ were exposed to two different concentrations of polystyrene beads: 1) 0.02 μg mL^-1^, and 2) 0.9 μg mL^-1^ of 0.3 μm (B0.3) and 1.1 μm (B1.1) beads, respectively. Those were incubated at 38.5°C in a humidified atmosphere of 5% CO_2_ and 95% O_2_ for 2 h, and 5 μL aliquots were taken at 0, 1, 1.5 and 2 h, were fixed with an equal volume of paraformaldehyde 4% and smeared across the slide. Those slides were left to dry at RT overnight before subsequent washes and analysis of bead attachment. Using the confocal Raman microscope, the slides were imaged with a ×100 DIC NA 0.9 objective, and random areas selected and analysed for the presence of sperm with microplastic attached to its surface. Each slide had at least 100 sperm cells counted (N = 4 bulls), and bead attachment was quantified.

For the other experiments, a total of 5x10^6^ sperm mL^-1^ were used for incubation, in which four treatment groups were prepared: 1) CT, a control group of RD media without beads; 2) bPS, RD media containing 0.7 μg mL^-1^ of PS beads; 3) esMP, RD media containing 0.369 μg mL^-1^ of PS beads; and 4) esPS, RD media containing 0.0257 μg mL^-1^ of PS beads. Sperm cells were incubated at 38.5°C in a humidified atmosphere of 5% CO_2_ and 95% O_2_ for 4 h, and assessed for motility, acrosome integrity, and oxidative stress at 0, 0.5, 1, 2, 3, and 4 h.

Following the same preparation procedure, 2x10^6^ sperm cells were exposed to the treatment groups already described (CT, bPS, esMP, esPS) for 1 h, incubated at 38.5°C in a humidified atmosphere of 5% CO_2_ and 95% O_2_. The sperm was then centrifuged (330xg, 5 min) and the pellet resuspended with 500 µL of RD media, this washing step was repeated two times for removal of polystyrene beads that are kept in suspension. After the second wash, the pellet was resuspended with 100 µL of fertilization medium (BO-IVF™, IVF Bioscience, UK), and used for *in vitro* fertilization (IVF). For each IVF round, an aliquot of the washed sperm was checked to exclude possible microplastics being transferred to the oocyte media,

### 2.8 Sperm motility, acrosome integrity and oxidative stress assessment

For the motility assessment, the mobile sperm analyzer iSperm® mCASA was used following the manufacturer’s instructions. Four fields of each sample were captured in each of the described time points, and the average motility used. The sperm progressivity was also analysed. For acrosome integrity and oxidative stress assessment, sperm cells were stained, at each time point, using NucBlue™ Live ReadyProbes™ Reagent (Hoechst 33342; for nuclear staining), 12.5 μM CellRox™ Green Reagent (Thermo Fisher Scientific, USA; for oxidative stress), and 5 μg mL^-1^ PNA-Alexa Fluor™ 647 Conjugate (for analysis of acrosome integrity), diluted in RD media. Aliquots of 5 μL sperm were taken, incubated for 15 min with the dyes solution, and fixed with 4% paraformaldehyde. Then, centrifuged at 1,700 rpm for 5 min, and the supernatant replaced by PBS. A second centrifugation allowed a wash for background removal. The pellets were finally resuspended in PBS and pipetted to a SuperFrost slide, smeared, and allowed to dry for at least 3 h. Then, the slides were washed 3 times for 5 min in PBS, mounted with antifade solution (ProLong™ Gold Antifade Mountant, Thermo Fisher Scientific, USA), and covered with a coverslip for posterior analysis, using an EVOS M7000 (Thermo Fisher Scientific, USA) microscope with a x20 NA 0.8 objective. Each slide had at least 100 sperm cells counted (N = 5 bulls).

### 2.9 *In vitro* embryo production

Bovine ovaries were obtained from a local slaughterhouse and the follicles between 2-8 mm were aspirated using an 18-gauge needle connected to a vacuum aspiration system. The precipitated pellets containing the cumulus-oocyte complexes (COCs) were transferred to a 9 cm Petri dish and 6 ml of wash medium (TCM 199 - GIBCO, buffered with 2.5% HEPES, supplemented with 10% FBS, 0.2 mM sodium pyruvate, 50 μg mL^-1^ gentamicin sulfate) previously warmed at 38.5°C was added. Around 200 grade I COCs were selected for each replicate, washed three more times in the same media, and transferred to maturation medium (TCM 199 - GIBCO, buffered with 25 mM sodium bicarbonate, supplemented with 10% FBS, 0.025 mM sodium pyruvate, 2.5 μg mL^-1^ gentamicin sulfate, 0.1 UI mL^-1^ recombinant human FSH) for a last wash before being divided into four wells (30-50 COCs per well) in a pre-equilibrate 4-well plate with 500 µl of maturation media at 38.5°C and 5% CO_2_ in humidified atmospheric air (21% O_2_) for 22 h.

Following, the COCs were washed using fertilization medium and transferred to new pre-equilibrated IVF-wells with 400 µl of fertilization medium in an incubator at 38.5°C and 5% CO_2_ in humidified atmospheric air (21% O_2_). The sperm was prepared as described above and 100 µL was added to each of the IVF-wells for incubation with the COCs at 38.5°C, 5% CO_2_, in humidified atmospheric air (21% O_2_) for 18 h.

Before starting the *in vitro* embryo culture (IVC), the 4-well plate was pre-equilibrated overnight in the incubator at 38.5°C, in a humidified atmosphere of 5% O_2_, 5% CO_2_, containing 500 μL of IVC media - ECS (previously described by (Santos et al., 2021)) and 400 μL of mineral oil (MP Biomedicals^TM^) per well. The presumptive zygotes were denuded by vortexing in a 15 ml tube with 1 ml of wash medium. Then, the denuded zygotes were washed two more times in the wash medium, one time in the embryo culture medium, transferred to the 4-well plate, and incubated at 38.5°C, in a humidified atmosphere of 5% O_2_, 5% CO_2_. Cleavage rate was evaluated on day 5 and, on day 7, the blastocyst rate was evaluated, and the blastocysts were stained for apoptosis and oxidative stress assessment.

### 2.10 Blastocysts apoptosis and oxidative stress assessment

To measure the levels of apoptosis and reactive oxygen species, the blastocysts were stained with NucBlue™ Live ReadyProbes™ Reagent (Hoechst 33342; for nuclear staining), 5 µM of CellRox™ deep red Reagent (Thermo Fisher Scientific, USA), and CellEvent™ Caspase 3/7 Detection Reagents in 300 µL of pre-equilibrated ECS media and incubated for 30 min at 38.5°C, 5% O_2_, and 5% CO_2_. The blastocysts were washed once in PBS containing 0.1% polyvinylpyrrolidone and imaged immediately after placing them on a glass slide. Images were taken on an EVOS M700 (Thermo Fisher Scientific). Each blastocyst was imaged individually with a 20x objective by taking Z-stacks with a 2 µm step size. The total number of cells and the apoptotic cells (caspase positive nuclei) were counted, and the apoptosis rate was determined as the percentage of caspase positive cells form the total counted cells. For oxidative stress, the maximum projection images were analysed using the ImageJ software by selecting the area of the embryo being imaged and quantifying the fluorescence intensity of the CellRox channel individually. The CellRox relative intensity was then determined by dividing the fluorescence intensity of the CellRox channel by the number of nuclei counted.

### 2.11 Statistical analysis

All functional data (i.e. sperm motility, acrosomal integrity, oxidative stress - sperm, cleavage and blastocyst rates, and embryo apoptosis) consisted of percentages that ranged between 0 and 100. We therefore assessed the effects of MPs on these functional traits via generalized linear regression models with binomial error distributions. For the sperm oxidative stress and bead attachment, however, we used a quasi-binomial error distribution to account for the fact that the data were under-dispersed. For embryo oxidative stress, the effects of MPs were assessed via generalized linear regression models with gaussian error distributions. In addition, we also applied a hierarchical approach in which data from each bull and replicate were allowed to have randomly varying intercepts. The significance was checked using a Tukey HSD. Differences were considered significant when p < 0.05. All data analysis and visualization were carried out in R (ver. 4.2.1).

## 3. Results

### 3.1 Microplastics and plastic related particles are present in seminal plasma from dogs and epididymal sperm from bulls

When searching for possible exposure concentrations of microplastics to perform biomimetic experiments, there is limited information in the literature. This is mainly because isolation of microplastics from complex biological samples is a challenge, which must be overcome and improved by the scientific community, especially when focusing on plastics of the nanometre size range. Therefore, our group had previously developed and optimized a protocol that was able to retrieve MPs from reproductive fluids samples (Grechi et al., 2023). Due to the complexity of the analysed matrix, and limited efficiency of digestion, many biological and big particles were recovered, increasing the analysis time of each sample for up to four or five days. The number of particles analysed ranged from 406 to 1,453, in canine seminal plasma sample 1 (SP1) and bovine epididymal sperm sample 1 (ES1), respectively. Only the particles that had a HQI (hit quality index) ≥ 75% were fully analysed, which represents 10% of the total.

Non-plastic related particles (NP) were present in all analysed samples, totalling an average 90.4 ± 72.5 particles mL^-1^ in dog seminal plasma (SP), and 186.7 ± 67.9 particles mL^-1^ in bull epididymal sperm (ES). Fibers (FB) were present in all the analysed samples, being the highest type of particles in SP and ES (128.8 ± 25.3, and 284.4 ± 362.9 particles mL^-1^, respectively). Other plastic related particles were also identified, such as coatings, solubilizers, and fillers (30.8, and 122.2 particles mL^-1^ in SP, and ES respectively), pigments (1.7, and 1.9 particles mL^-1^ in SP, and ES respectively), and plasticizers (70.8, and 170.3 particles mL^-1^ in SP, and ES respectively). In total, 43 different polymers were identified in all the samples, and the most abundant were polypropylene (25 particles mL^-1^ in ES3), cellulose (13.3 particles mL^-1^ in ES2), silicon(IV) oxide (9.2 particles mL^-1^ in SP2, 15 particles mL^-1^ in ES1), and polystyrene (6.7 particles mL^-1^ in SP1). Polystyrene and silicon(IV) oxide were the most abundant plastic particles in SP, and polypropylene in ES samples (Figure 1a). Considering the aimed exposure concentration for the following experiments, it was calculated the weight of total microplastics and the weight of polystyrene found in the epididymal sperm samples, based on the size and density of the plastic particles found. The average concentration of MPs found in ES was 0.37 µg mL^-1^. In ES1 no polystyrene particles were identified, and the average PS weight in all bovine epididymal sperm was 0.026 µg mL^-1^. The MPs found in SP had an average of 14.7 ± 12.8 µm in length and 6.6 ± 4.5 µm in width. In ES the size average was 18.4 ± 25.4 µm in length and 8.5 ± 8.8 µm in width (Figure 1b and c).

**Figure 1.**
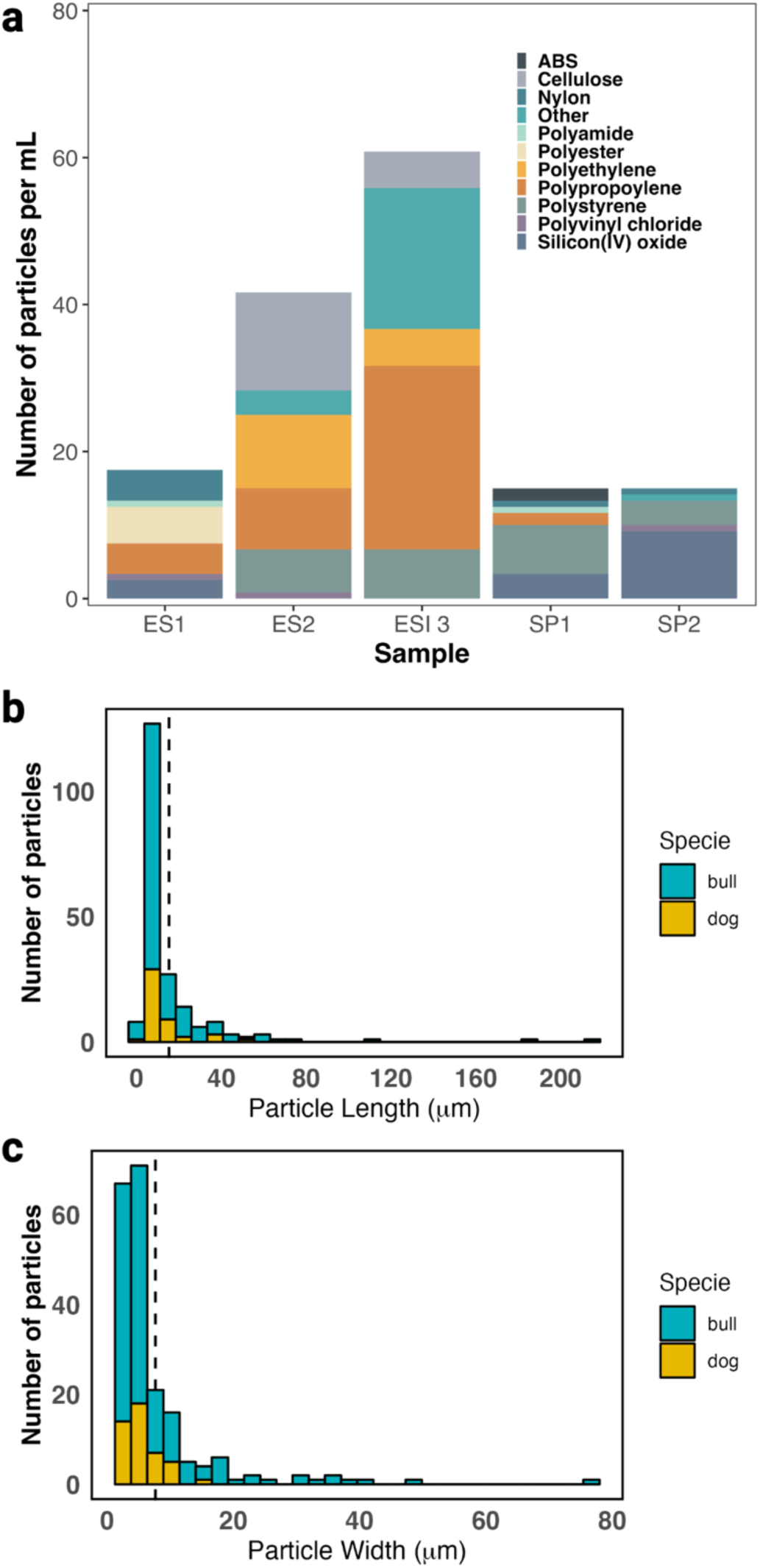
MPs were detected in bovine epididymal sperm (ES), and canine seminal plasma (SP). In (a) the polymer distribution of the particles found in the samples by Raman spectroscopy. In (b) and (c) the length ns width distribution of the found polymers is displayed, respectively. Dashed lines in (b) and (c) represent the average length and width of the found particles in both bull and dog samples (16.5 and 7.55 µm, respectively).

### 3.2 Polystyrene MPs bind to the sperm surface and affect sperm motility

To investigate possible variances between the different sizes of MPs on the attachment and hypothesizing a possible internalization of MPs, the smaller (0.3 µm) and the bigger (1.1 µm) beads were chosen to be analysed for sperm attachment. Sperm incubated in the presence of the 0.3 μm beads had a higher number of beads attached to the surface than those incubated with the 1.1 μm beads (p < 0.001). We also found that the percentage of sperm with attached MP beads did not change over time (p = 0.3), suggesting that attachment occurs very early on after exposure and may not be reversible (Figure 2a). From the analyses, we found no evidence of polystyrene beads being internalized by the sperm cells.

**Figure 2.**
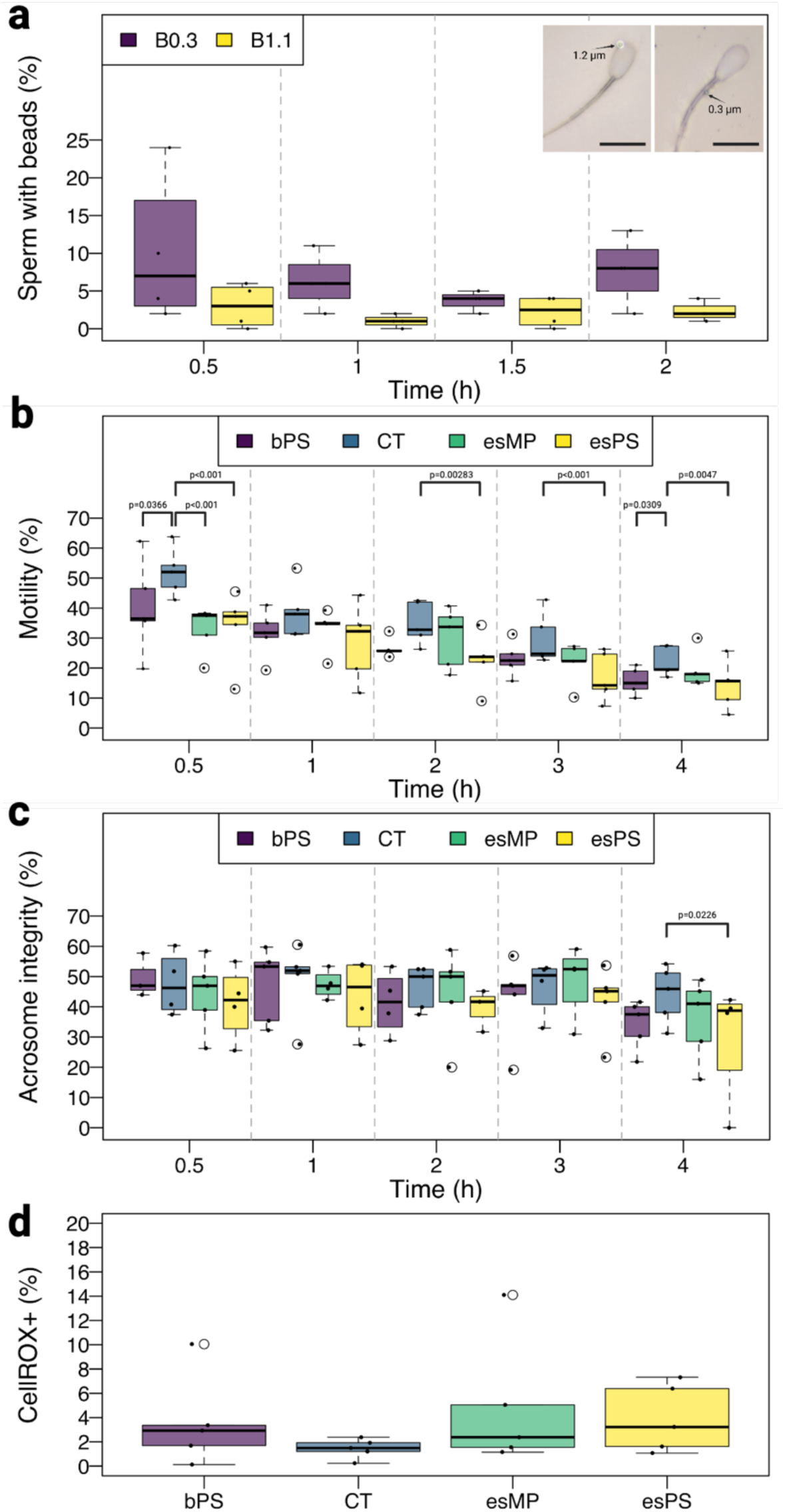
PS beads alter bovine sperm parameters in short-term exposure. Sperm exposed to PS do not internalize beads, and 0.3 µm beads are able to attach more to the sperm surface than 1.1 µm beads, and is not time dependent (a). In (b) the sperm motility is affected in all groups in the first analysed time point (0.5 h), and lowest concentration exposure to MPs maintains effects over time. PS was not able to affect sperm acrosome integrity (c) or ROS production (d). Scale bar = 10 µm.

As normally expected for cryopreserved sperm, the sperm motility decreased over time (p < 0.001). We also observed that PS beads significantly decreased sperm motility at time point 0.5 h compared to the control (p = 0.0366, p < 0.001, and p < 0.001, for bPS, esMP, and esPS, respectively). The group exposed to the smallest concentration of polystyrene (esPS) also showed a statistical difference at time points 2, 3, and 4 h (p < 0.01) when compared to the control group (Figure 2b), in addition to the highest concentration (bPS) also showing differences after 4 h of exposure (p = 0.0309). No significant differences were found in the progressivity of sperm cells (Supplementary file S2). In addition to the motility assays, acrosome integrity (Figure 2c) and oxidative stress (Figure 2d) of the sperm cells exposed to PS were assessed. MP polystyrene beads did not affect acrosome integrity, nor increased production of ROS in sperm exposed to true to life concentrations. Although no statistical difference was observed for oxidative stress, it is possible to see that when compared to the control, the group esMP showed a trend to have increased oxidative stress (p = 0.0567). In conclusion, a mix of different sizes of PS beads in true to life concentrations was able to affect sperm motility in a short-term exposure, but not sustain the effect over time, and did not affect acrosome integrity or oxidative stress.

### 3.3 Oocytes fertilized with sperm exposed to MPs had reduced developmental potential and resulting embryos had increased oxidative stress and apoptosis rates

To evaluate the potential impact of PS MPs exposure on *in vitro* fertilization outcomes, we used sperm samples exposed to varying MP concentrations. No statistically significant differences in cleavage rates were observed across the experimental groups when compared to the control group (Figure 3a). However, when analyzing embryo development on day 7, although no differences on expanded and hatching blastocysts were observed, the group exposed to the highest MP concentration (bPS) exhibited a significant reduction on blastocyst rate compared to the control group (29.8 ± 4.4 vs 19 ± 7.7%, respectively; p = 0.0344) (Figure 3b). In addition to assessing blastocyst formation, we also investigated oxidative stress and apoptosis levels in the embryos. Remarkably, all groups that were exposed to sperm previously in contact with MPs exhibited a substantial increase in ROS formation when contrasted with the control group (p < 0.001; Figure 3c). Blastocyst formed from bPS exposed sperm also displayed a significant increase in the number of apoptotic cells (p < 0.001) (Figure 3d). While the other groups did not demonstrate statistically significant differences, a trend in having higher apoptosis was observed across all groups in comparison to the control (CT). This observation suggests a potential association with the MP concentrations tested. Specifically, the control group exhibited an apoptotic cell rate of 3.9 ± 3.9%. In contrast, the bPS group had 9.3 ± 5.8 (p < 0.001), esMP 7.2 ± 5% (p = 0.0284), and esPS 6.8 ± 6% (p = 0.0643). Figure 3e shows examples of apoptosis and ROS staining of the different groups.

**Figure 3.**
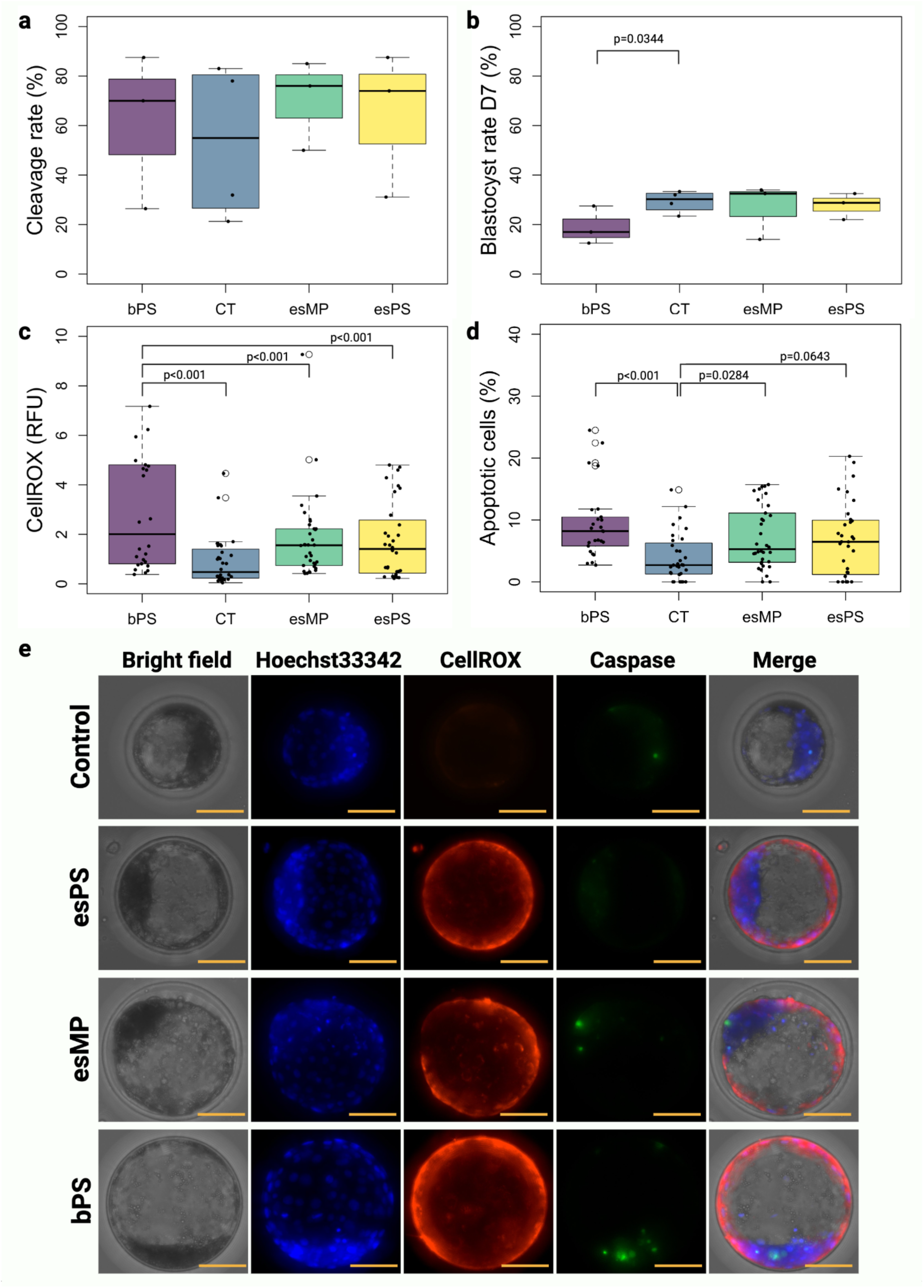
Sperm exposed to PS previously to fertilization can influence oxidative stress and apoptosis of embryos. After fertilization, no effects of PS are displayed in the cleavage rate (**a**) between the control (CT, N=31), blood PS (bPS, N=34), epidydimal sperm MP (esMP, N=34) and epidydimal sperm PS (esPS, N=32), but the highest exposure concentration of PS (bPS) was able to decrease the blastocyst rate on day 7 (**b**). Sperm exposed to PS MP increased ROS production (**c**), and apoptosis rates (**d**) in the resulting embryos. In (**e**) example fluorescence images of apoptosis and oxidative stress of embryos are shown. Scale bar = 50 µm.

To ensure that there were no observable effects of any residual MPs during the sperm wash process when introduced into the fertilization media, we conducted a pilot study. In this study, MPs were processed in the same manner as described previously for sperm MPs’ incubation and preparation for IVF, but without the presence of actual sperm. Subsequently, the same volume of these prepared MPs was added to the IVF well, followed by fertilization using sperm that had not been incubated with MPs. The purpose of this pilot study was to ascertain whether any effects observed in the developing embryos could be attributed to the potential transfer of MPs into the IVF well, possibly due to residual MPs remaining after the sperm wash process. Our findings revealed no significant differences between the control group and the group exposed to MPs without sperm during the wash process concerning embryo cleavage, blastocyst formation rate, apoptosis, and oxidative stress. These results suggest that the observed effects on embryos are primarily a consequence of the influence of MPs on spermatozoa, rather than any remaining MPs in the fertilization media.

## 1. Discussion

Microplastics are small plastic particles distributed all over the world, from remote mountains to deep-sea sediments, even making their way into air, soil, water, and the food chain. Recent studies show that these MPs have also entered the body, being found in different human tissues, which emphasizes the need to understand their impact on people, animals, and the environment. The rise of single-use plastics in our daily lives has coincided with a drop in fertility rates, which raises questions about whether there’s a link between microplastics and infertility. While preliminary connections have emerged (Zhang et al., 2022), *in vivo* studies have often used unrealistically high concentrations of MPs, creating an incongruity between laboratory investigations and real-world exposure scenarios (Mills et al., 2023). To overcome this, we carefully optimized a method previously developed by our group to measure MP particles in bull sperm and dog semen (Grechi et al., 2023). With these results, we were able to expose sperm to realistic levels of MPs to see how they affect sperm and the outcomes of *in vitro* fertilization.

Isolating MPs from complex biological materials remains a big challenge. In our study, we were able to isolate several particles from bovine epididymal sperm and canine semen, showing that MPs can reach those compartments. Nevertheless, we were limited to MPs bigger than 1 µm, due to limitations of the digestion protocol, and the detection limit of the microscope used for the analyses. Our analysis revealed MP presence in all seminal plasma and epididymal sperm samples from bulls and dogs. This detection is consistent with human studies where MPs were detected in a substantial proportion of samples, specifically 6 out of 10 (Montano et al., 2023) and 11 out of 25 (Zhao et al., 2023b) samples analysed. The lipophilicity of MPs suggests a natural affinity for lipid-rich tissues, such as the testis and epididymis, potentially facilitating their transfer into semen. Supporting this is a study by (Sen et al., 2022), where mice were exposed to fluorescent polystyrene beads of 0.1, 3, and 10 μm. Post-exposure, MPs were detected in the testis and epididymis, with the latter showing notably higher levels. This accumulation could explain the MP presence in semen. The study also indicated that the transport of MPs within the body is time and size-dependent, with smaller beads being detected more readily than larger ones. These findings, mirrored by our observations in animal reproductive tissues, suggest a similar pattern of MP behaviour across mammalian species and underscore the importance of understanding MP dynamics in reproductive health.

Here, out of the 43 distinct polymers identified, the most abundant were PP, cellulose, silicon, and PS. The average dimensions of MPs in SP (14.7 µm in length and 6.6 µm in width) were smaller than in ES (18.4 µm in length and 8.5 µm in width) samples. Compared to a human study, where microplastics were smaller, ranging from 2 to 6 µm, our findings indicate a broader size distribution in animal samples. This study primarily reported irregularly shaped fragments, with the most common polymers being those frequently found in everyday items such as PP, PS, PET, PE, POM, PVC, and PC, with some samples even containing acrylic micro fragments (Montano et al., 2023). Particularly notable is the study by (Zhao et al., 2023b), which reported much larger particles, with an average size between 20 to 100 μm in semen and testis, predominantly composed of PVC and PE in various shapes in semen, and PS fragments in the testis. The average abundance reported was 0.23 ± 0.45 particles mL^-1^ in semen and 11.60 ± 15.52 particles g^-1^ in testis. In stark contrast, the MPs detected in our animal samples, although varied in polymer type, were generally smaller, especially in seminal plasma, highlighting a potential difference in MP profiles between species or in the methodologies used for MP detection and quantification.

After creating a method to find realistic plastic levels and being able to overcome the problem of overestimation for exposure experiments, we investigated how such particles could have an effect in reproductive parameters, so we looked at how MPs might affect bull sperm. Similarly to mice and rats fed with PS MPs (Hou et al., 2021; Ijaz et al., 2021; Jin et al., 2022, 2021; Li et al., 2021), we saw varying effects of MPs on sperm motility, being reduced in the esPS and the bPS exposed groups at different time points. In humans, there is evidence suggesting that seminal samples containing MPs are correlated with lower semen quality in terms of volume, sperm number, motility and morphology (Montano et al., 2023). A more evident change in sperm motility was seen in mice and rats after *in vivo* exposure to PS MPs (Ijaz et al., 2021; Jin et al., 2022, 2021). Here, however, frozen-thawed sperm cells experienced a direct 4 h exposure to 0.0257 - 0.7 μg mL^-1^ PS, whereas in *in vivo* studies mice were exposed to 1,000 μg mL^-1^ day^-1^ for 28 days, and rats to 15 – 1,500 μg day^-1^ for 90 days. Differently from existing studies, we did not observe reduced acrosome integrity and increased oxidative stress in sperm exposed to PS MPs. However, research with mice showed that feeding them PS, led to oxidative stress, especially in those fed amounts ranging from 2 to 2,000 μg mL^-1^ (Ijaz et al., 2021). Similarly, another study that administered PS to mice evidenced disruptions in acrosome integrity for doses of 1 and 10 mg kg^-1^ (Zhou et al., 2022). It is important to note that such researchers studied the effects of MPs on male fertility by feeding mice MPs at concentrations that were ca. 100,000 times greater than they would be exposed to in the wild (Mills et al., 2022). Our findings, in contrast, have exposed sperm directly to a low concentration of PS MPs for a short period of time. The reduced acrosome integrity in such studies can be directly associated with the high concentrations of exposure, in addition to effects that can be associated with spermatogenesis, which were not able to be tested in this study. While these observations in humans, mice and rats provide a concerning parallel to our findings, the direct effects of MPs on sperm health across species necessitate further investigation to understand the implications fully. It’s important to note that our *in vitro* exposure model might not capture the complexities of *in vivo* conditions, such as bioaccumulation, systemic distribution, and long-term tissue effects of MPs, which could exacerbate the impact on sperm motility and overall seminal quality.

Remarkably, the incubation of sperm with PS MPs, while not significantly affecting the sperm parameters measured in our study, had profound implications during IVF. The use of PS-exposed sperm in IVF resulted not only in embryos with heightened levels of apoptosis and oxidative stress but also in reduced blastocyst development. This suggests that PS microplastics might exert sublethal influences on sperm which, though not detectable in standard sperm assessments, could lead to detrimental outcomes during early embryonic development. There is substantial evidence of the adverse effects of MPs on embryo development in a range of species. For example, different studies indicate MPs affect marine copepods, monogonont rotifers, and Pacific oysters, decreasing egg size and hatching rates, offspring numbers, and larval development (Cole and Galloway, 2015; Jeong et al., 2016; Sussarellu et al., 2016). Similarly, invertebrates like *Hyalella azteca* and aquatic crustaceans such as *Ceriodaphnia dubia*, as well as terrestrial worms and springtails, experience reduced neonates and juvenile numbers due to MPs (Au et al., 2015; Ju et al., 2019; Lahive et al., 2019; Zhu et al., 2018; Ziajahromi et al., 2017). Nematode worms also show diminished reproductive potential, including decreased embryo numbers (Lei et al., 2018; Schöpfer et al., 2020).

In light of these findings, our research extends the current understanding by providing novel evidence that MPs can affect sperm quality, which subsequently has a negative impact on embryo development. This link between MP exposure, compromised sperm function, and subsequent embryonic development has not been previously demonstrated, marking our study as a pivotal contribution to the field. While not statistically significant, sperm exposed to MPs exhibited an increase in oxidative stress, a phenomenon seemingly transmitted to developing embryos. Previous studies have demonstrated that sperm subjected to oxidative stress display compromised traits such as decreased motility, pro-oxidation, and premature capacitation, all of which can directly impact fertilization (de Castro et al., 2016). Even at low levels of oxidative stress that may go unnoticed, sperm have the capacity to carry damaged DNA, which can adversely affect the health of the resulting offspring (Aitken and Baker, 2006; Gavriliouk and Aitken, 2015). As seen here, it is widely acknowledged that uncorrected damage in sperm may not impede fertilization capability but may only manifest effects during embryo development (Bollwein and Bittner, 2018).

While there aren’t many studies on how MPs affect fertility, the ones that exist highlight their importance as harmful environmental pollutants. Here, we demonstrated that MPs can reach both bull epidydimal sperm and dog seminal plasma. Moreover, we have showed, for the first time, how exposing sperm to a mix of different-sized PS particles in biological found concentrations, can have consequences for embryo development after fertilization, reinforcing the urgency for research and strategies to address the potential threat those particles pose to human and animal fertility.

## Abbreviations

ρ: Particle density
bPS: Blood polystyrene
COCs: Cumulus-oocyte complexes
CSF: Coatings, solvents, or fillers
ES: Epididymal sperm
esMP: Epididymal sperm total microplastics
esPS: Epididymal sperm total polystyrene
FB: Fibers
IVC: *In vitro* embryo culture
IVF: *In vitro* fertilization
MP(s): Microplastic(s)
NP: Non-plastic related particles
PFA: Perfluoroalkoxy alkane
PG: Pigments
PP: Plastic polymers
PTFE: Polytetrafluoroethylene polymer
Pw: Plastic weight
PZ: Plasticizers
ROS: Reactive oxygen species
RT: Room temperature
SP: Seminal plasma
UK: Unknown
v: Particle volume

## CRediT authorship contribution statement

Conceptualization: MAMMF; Methodology: MAMMMF, NG, GF, SD; Investigation: NG, GF and SD; Supervision: MAMMF; Writing—original draft: NG and MAMMF; Writing—review & editing: all authors.

## Supplementary data

Supplementary file S1: Supplementary Figure S1.

## Data availability

All data are available in the main text or as supplementary data.

