## Supplementary File S1 for "Microplastics are detected in bull and dog sperm and polystyrene microparticles impair sperm fertilization"

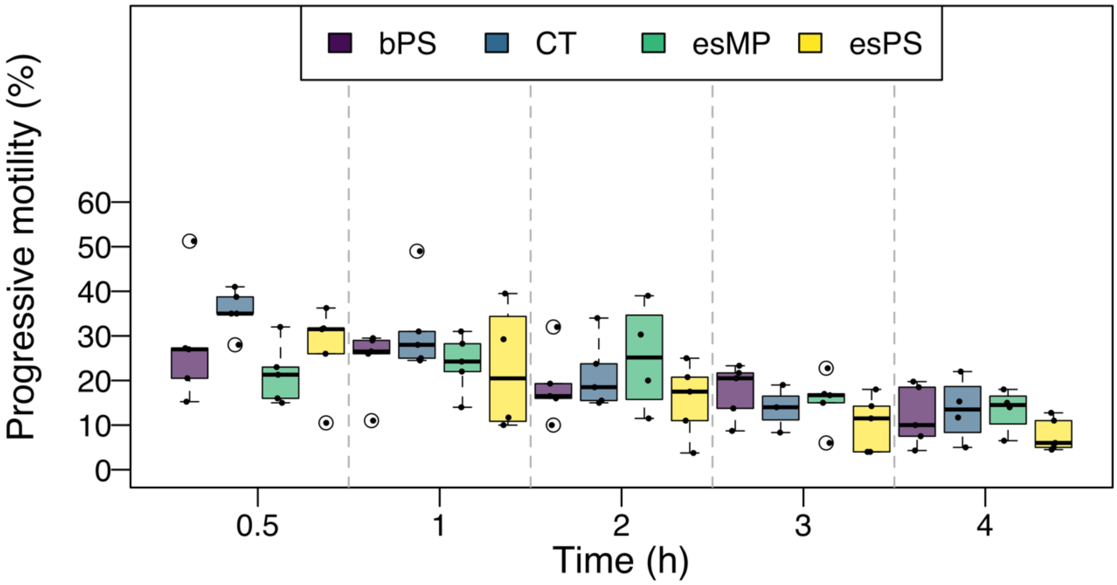


**Supplementary Figure S1.** PS beads do not alter progressive motility of bovine sperm in short-term exposure.
